# Intraspecific genetic variation is critical to robust toxicological predictions of aquatic contaminants

**DOI:** 10.1101/2023.06.06.543817

**Authors:** René S. Shahmohamadloo, Seth M. Rudman, Catherine I. Clare, Judy A. Westrick, Xueqi Wang, Luc De Meester, John M. Fryxell

**Author notes:** These authors contributed equally to this work (René S. Shahmohamadloo, Seth M. Rudman). Address co-correspondence to: René S. Shahmohamadloo, PhD, Liber Ero Postdoctoral Fellow, NSERC Postdoctoral Fellow School of Biological Sciences | Washington State University, 14204 NE Salmon Creek Avenue | Vancouver | Washington | 98686 | United States. Address co-correspondence to: Seth M. Rudman, PhD Assistant Professor, School of Biological Sciences | Washington State University, 14204 NE Salmon Creek Avenue | Vancouver | Washington | 98686 | United States.

## Abstract

Environmental risk assessment is a critical tool for protecting aquatic life and its effectiveness is predicated on predicting how natural populations respond to contaminants. Yet, routine toxicity testing typically examines only one genotype, which may render risk assessments inaccurate as populations are most often composed of genetically distinct individuals. To determine the importance of intraspecific variation in the translation of toxicity testing to populations, we quantified the magnitude of genetic variation within 20 *Daphnia magna* clones derived from one lake using whole genome sequencing and phenotypic assays. We repeated these assays across two exposure levels of microcystins, a cosmopolitan and lethal aquatic contaminant produced by harmful algal blooms. We found considerable intraspecific genetic variation in survival, growth, and reproduction, which was amplified by microcystins exposure. Finally, using simulations we demonstrate that the common practice of employing a single genotype to calculate toxicity tolerance failed to produce an estimate within the 95% confidence interval over half of the time. These results illuminate the importance of incorporating intraspecific genetic variation into toxicity testing to reliably predict how natural populations will respond to aquatic contaminants.

## Main Text

Reliable translation of aquatic toxicity testing into predictions of population-level consequences in nature is a cornerstone of environmental risk assessments ^1–6^. Such predictions are classically obtained by quantifying individual phenotypic responses under constant conditions in a laboratory ^7^. Research has historically favored single-species toxicity testing that generates point estimates such as the lethal concentration at which a contaminant will kill 50% of a test population (LC50) ^8^. For more than fifty years, regulatory agencies have relied on such lethality data to inform environmental risk assessments, mandated to safeguard biodiversity and associated ecosystem services by protecting rare or economically significant species ^6^. Yet, reliably translating laboratory toxicity studies into population-level predictions requires building on traditional methods to incorporate aspects that are key to ecological realism ^8^.

Genetic variation within species is recognized as a critical driver of both ecological and evolutionary processes ^9–11^. In ecological contexts, genetic variation begets variation in functional traits, which can have profound impacts on populations, communities, and ecosystems ^9–12^. Genetic variation within species is central to evolution, as the process of natural selection shaping genetic variation is what produces adaptation ^10,13^. Among the most visible and societally challenging examples of rapid adaptation are cases where populations adapt to agrochemicals ^14^ or contaminants ^15^. Despite an appreciation of the role of adaptation in resistance and some demonstrations of the amount of standing variation in tolerance ^4,5,16–18^, recognition of the importance of intraspecific genetic variation continues to be ignored by regulators within the discipline of toxicology and, more broadly, the environmental risk assessment process.

Investigating the magnitude of intraspecific variation within an aquatic population represents a crucial and understudied element in toxicology and more broadly environmental monitoring programs. Little is known about the magnitude and ecological importance of intraspecific genetic variation in toxicological responses within natural populations. Choosing a genotype based on availability (e.g., standard lab strain) may lead to inferences that are unrepresentative of imperiled populations. More broadly, conducting measurements on any single genotype is using a single point to estimate a distribution that likely has substantial variance ^19,20^. Detailed study of specific populations that includes both measures of the extent of genetic variation within focal populations and classical toxicity testing are needed to infer the potential for this variation to impact population-level responses under toxic insults ^21^.

*Daphnia magna* (water fleas) are a powerful eco-genomic model system because they are simple to culture, have a fully-sequenced genome, and they play an important and well-documented ecological role in many freshwater ecosystems ^22,23^. With short generation times and considerable genetic diversity derived from diapausing eggs in sediments, *Daphnia* can undergo rapid adaptation in response to environmental variation ^22–26^. These eco-evolutionary insights have not translated to change in routine laboratory testing of *Daphnia*, which is a model system used in toxicology to test chemical compounds before they are produced, released or sold ^23^. Toxicologists previously advocated that toxicological studies should account for genetic variation in *Daphnia* based on discrepancies between several laboratory strains that were exposed to the same reference toxicants ^2,3,7,27^. Yet, determining the loss in accuracy associated with choosing a single clone from a natural and genetically heterogenous population for environmental risks assessments is crucial to demonstrating the overall importance of accounting for genetic variation.

Determining the magnitude of variation within a population is crucial both for predicting immediate toxicological responses and as an indicator of the potential for rapid adaptation. Doing so requires accounting for the scale and scope of genetic diversity within a population and conducting toxicity testing on a heterogeneous and realistic population of *Daphnia*. Genomic data can be useful in meeting these objectives as genome-wide data provide information on the scale and scope of genetic diversity being studied and can be translated to provide population-level insight. For investigations on the relationship between genetic diversity and toxicological responses whole genome sequencing data yields an ability to identify populations or clones and compare the extent to which they are genetically divergent, to determine whether the degree of genomic differentiation between populations is correlated with toxicological responses, and to test for relationships between genomic diversity and tolerance. Depending on the underlying genetic architecture of variation in toxin tolerance it may be possible to identify the genetic basis of resistance ^28^, though finding a ‘simple’ genetic basis may not be common ^29^. The use of candidate loci, identified through prior mapping efforts or through changes in gene expression, provides an opportunity to test hypotheses about the relationship between genetic variation in potentially important genes and toxicological responses that could link genetic and phenotypic variation ^30–32^. At a minimum, incorporating population genetic information can help standardized toxicity tests and the information generated also has the potential to better predict the risk of toxicant exposure on populations, and increase predictability on the associated community and ecosystem functions.

To determine the importance of intraspecific variation in toxicity projections key to the environmental risk assessment process, we sequenced the genomes of 20 *Daphnia magna* clones collected from a single Belgian lake and quantified the chronic effects of the cyanobacterium *Microcystis* on their life-history traits (survival, growth, reproduction, broods). Harmful algal blooms (HABs) of *Microcystis* are a prominent and toxic ecological disturbance ^33^ that strongly impact a wide-range of aquatic taxa due to the production of microcystin toxins ^34–39^. As a keystone zooplankton grazer, *Daphnia* shows substantial intraspecific diversity and rapid adaptation in response to HABs ^24,40,41^, including some evidence that particular genotypes can effectively feed on and reduce the abundance of toxic *Microcystis* ^42,43^. Determining whether intraspecific variation in toxin tolerance influences vital demographic rates and population dynamics by *Daphnia* clarifies the role of intraspecific genetic variation in toxicology, with implications for regulatory agencies tasked with maintaining biodiversity under toxic bloom conditions.

In this study, 30 individuals per genotype were divided and exposed to three treatments (i.e., common gardens): 100% green algae (*Chlorella vulgaris*, non-toxic diet), 2:1 chlorella:microcystis (a moderately toxic diet), and 1:1 chlorella:microcystis (a severely toxic diet). Common garden rearing keeps environmental conditions constant allowing for the estimation of the contribution of genetic variation to phenotypic variation. We chose 2:1 and 1:1 toxic treatments that resulted in microcystin concentrations of 3.3 ± 0.001 μg L^−1^ and 5.1 ± 0.001 μg L^−1^ (Dataset S1), respectively, which have been considered sublethal ^44^ to lethal ^45,46^ in *Daphnia* laboratory assays and are common ratios in freshwater ecosystems with HABs ^36^. We sought to address the following four questions: 1) How much intraspecific genomic and phenotypic variation occurs within a single population of *D. magna*? 2) Does exposure to toxic cyanobacteria reduce the magnitude of intraspecific phenotypic variation? 3) Is there an interaction between clonal growth and reproduction under favorable (green algae) and toxic (cyanobacteria) conditions? 4) Using data from these trials, how would sampling design influence the interpretation of reproductive output with and without contaminant exposure? We predicted that *D. magna* clones would exhibit considerable genetic and phenotypic intraspecific variation in toxicological responses, as *Daphnia* populations have been shown to be variable. Additionally, we predicted that phenotypic variation would be reduced when exposed to a toxic insult due to a decrease in trait values in high performing genotypes. Moreover, we predicted a trade-off between growth and reproduction under favorable and toxic conditions, with clones having the greatest reproductive output varying across cyanobacterial concentrations. Finally, we predicted that sampling a sufficient number of genotypes would be critical to producing an accurate estimate of reproduction in both favorable and toxic conditions.

### Intraspecific genomic and phenotypic variation

Genetic variation among *D. magna* clones was considerable in both phenotype and genotype (Figure 1). When reared on a *Chlorella*-only non-toxic diet, phenotypic variation among clones was significant for survival (*Χ*^2^ = 4.52, *P* = 0.033), growth (*Χ*^2^ = 67.40, *P* < 0.0001), and reproduction (*Χ*^2^ = 40.22, *P* < 0.0001). The magnitude of genome-wide divergence between pairwise sets of clones varied (Figure 1a), with the most similar clones having ~1.2M SNP differences and the most divergent ~1.6M.

**Figure 1.**
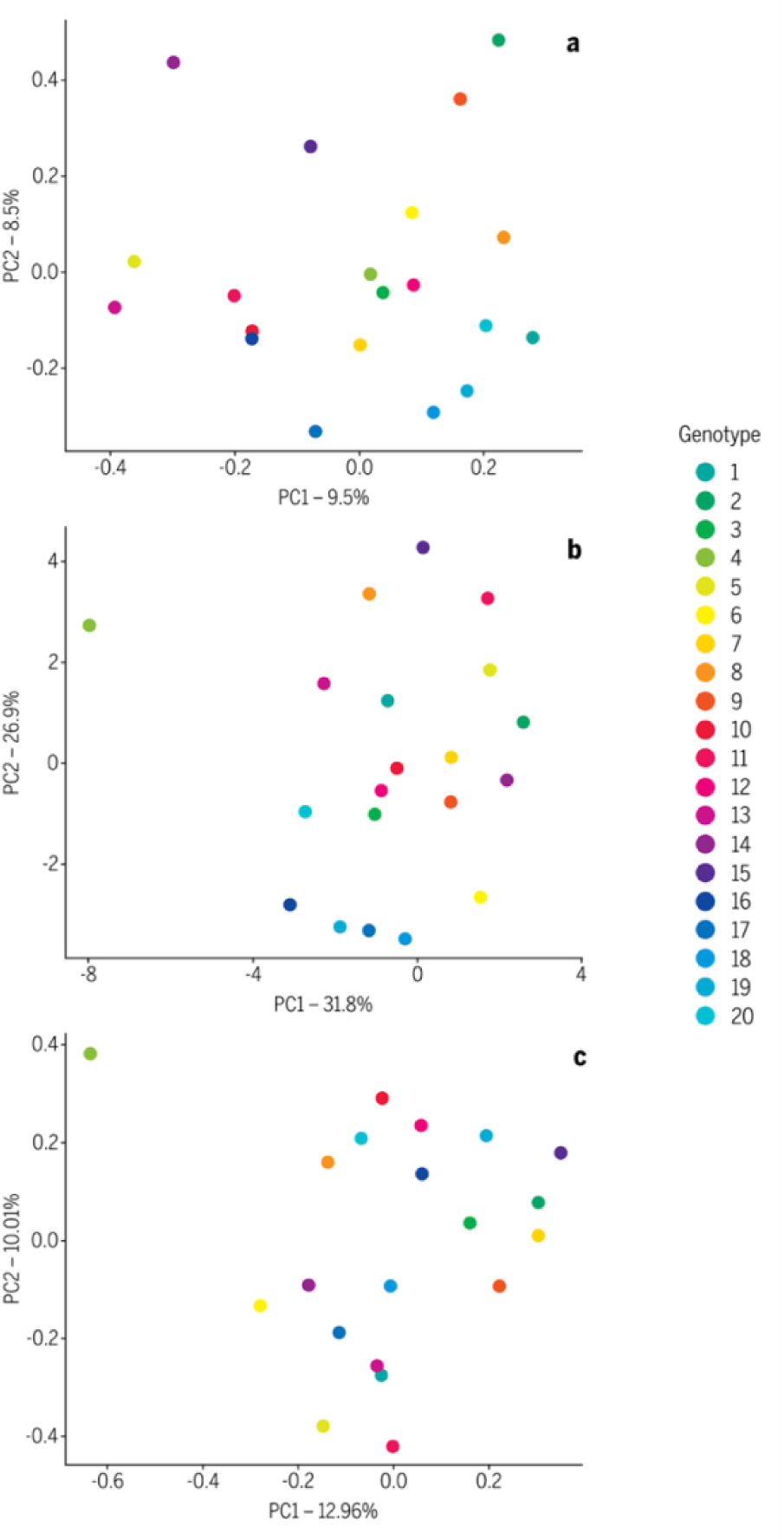
Multivariate a) whole genome genomic and b) phenotypic divergence across the all treatments between 20 *Daphnia magna* clones visualized using principal component analysis (PCA). Panel c) shows a PCA constructed from genetic variation at putative candidate loci for variation in tolerance to *Microcystis* in *Daphnia*.

We tested for an association between the degree of genomic divergence between clones and the magnitude of their phenotypic differentiation. We calculated a phenotypic distance between each pair of clones based on results of phenotypic assays for survival, growth, time to reproduction, and total reproduction when reared on non-toxic, moderately toxic, and severely toxic diets. We found no significant association between the degree of genomic divergence genome-wide and the magnitude of overall phenotypic divergence between clones (Figure 1) (mantel *r* = 0.033, *P* = 0.781). We also examined the relationship between genetic diversity found within clones (i.e., total heterozygosity) and clonal survival and reproduction (Supporting Information, Dataset S1). We found no significant association between the amount of genetic diversity within clones and their ability to survive or reproduce across diet conditions.

We used a similar analysis approach to assess the association between phenotypic differentiation between clones and genetic variation in 8 gene regions previously implicated in *Daphnia* transcriptomic responses to microcystins ^47,48^. A PCA of the all variant sites with these 8 gene regions revealed that clone 4, a phenotypically distinct clone due to its overall low survival and growth rates across conditions (Figure 1B), is largely separated from all other clones by PC1. We found no significant association between the amount of divergence at these putatively functional gene regions and the magnitude of overall phenotypic divergence between clones (Figure 1) (mantel *r* = 0.054, *P* = 0.584). We also found no association between the amount of genetic diversity at these putatively functional gene regions and their ability to survive or reproduce at high (1:1) *Microcystis* exposure.

### Magnitude of phenotypic variation from toxic exposure

Exposure to cyanobacteria reduced the overall fitness of *D. magna* genotypes following a monotonic dose response in the moderately toxic (2:1 *Chlorella* : *M. aeruginosa*) and severely toxic (1:1 *Chlorella* : *M. aeruginosa*) treatments (Figure 2). For example, mean survival across all clones for the 14-day trial declined from 94% in the non-toxic diet, to 86.5% in the moderately toxic diet, and to 53% in the severely toxic diet. We tested for differences in the magnitude of clonal variation with increasing exposure to *M. aeruginosa* for each of the three fitness-associated phenotypes (survival, growth, reproduction). Intraspecific variation in survival did not increase from the control to 2:1 (*F\** = 3.60, *P* = 0.066) but did from 2:1 to 1:1 (*F*\* = 22.59, *P* < 0.001). Intraspecific variation in growth rate increased from the control to 2:1 (*F*\* = 10.28, *p* = 0.003) but decreased from 2:1 to 1:1 (*F*\* = 11.82, *P* = 0.002). Neonate production showed the most consistent pattern, with intraspecific variation between clones declining with increasing *M. aeruginosa* exposure (control to 2:1 (*F*\* = 18.54, *P* = 0.0001); 2:1 to 1:1 (*F*\* = 7.44, *P* = 0.009)).

**Figure 2.**
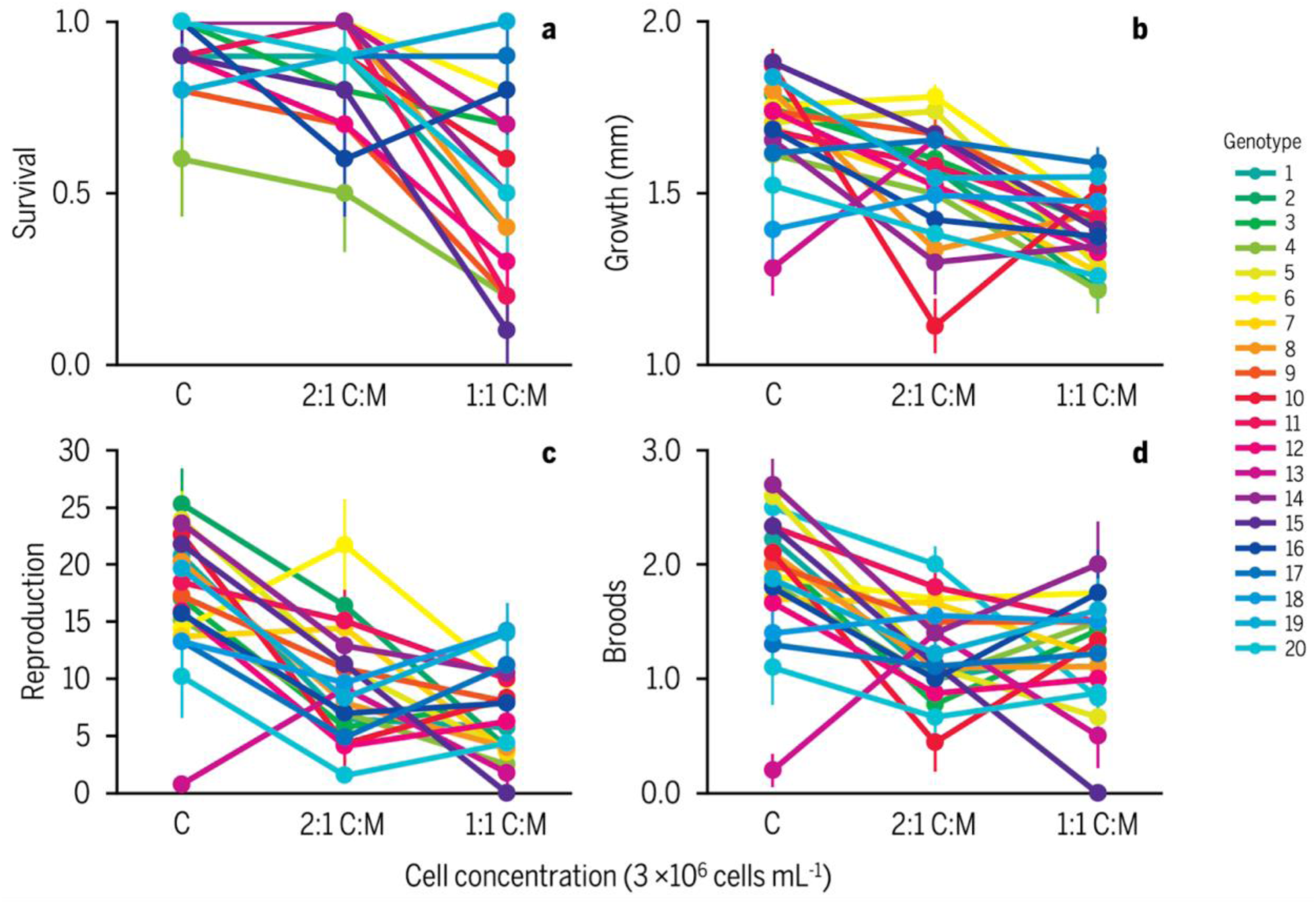
Phenotypic variation among the 20 *Daphnia magna* clones at the end of the 14-d chronic toxicity study. Phenotypes measured in parents were a) survival, b) growth from day 0 to 14, c) total number of neonates produced, and d) total number of broods produced. Cell concentration was standardized across three algal food treatments: non-toxic - “C”, moderately-toxic - “2:1 C:M”, and severely-toxic - “1:1 C:M”. Plots show means and standard errors for each clone when reared on each diet.

### Interactions under favorable and toxic conditions

We tested for interactions associated with performance across variations in diet toxicity. On average, survival declined with increasing toxicity by 8% in the moderately-toxic diet and 44% in the severely toxic diet relative to controls. However, we found evidence of an interaction between clonal identity and a *Microcystis* diet (*Χ*^2^ = 59.68, *P* = 0.014). Some of these interactive effects were striking. Relative to rearing on the *Chlorella* diet, clones 5 and 15 showed an 80% decline in mean survival when reared on the severely toxic diet while clone 19 exhibited a 20% increase in survival. Similarly, we found a significant interaction associated with growth across treatments (*Χ*^2^ = 244.17, *P* < 0.0001). Finally, neonate production showed a similar pattern, with a significant interaction between clonal identity and a *Microcystis* diet (*Χ*^2^ = 264.42, *P* < 0.0001).

Of particular interest was that certain genotypes performed better in one of the toxic treatments compared to the *Chlorella* diet (e.g., genotypes 6 in 2:1, and 13 in both 2:1 and 1:1, across all life-history traits), while other genotypes performed better in the 1:1 versus 2:1 toxic treatments (e.g., genotypes 10, 14, 18, 19, 20). These findings indicate that relative performance of clones is dependent on the magnitude of toxicant exposure. reinforce the shifts in relative performance across clones associated with changing *M. aeruginosa* conditions.

### Influence of genetic diversity on toxicological inferences

We used a bootstrapping approach to determine how variation in the number of clones in the experimental sampling design could influence the robustness of the data relative to including 20 clones. Our results demonstrate that sampling individuals from a larger number of genotypes is critical to generating a robust estimate of reproduction in all 3 rearing conditions (Figure 3a). To quantify the loss of robustness and accuracy that would result from assaying a reduced amount of genetic diversity we assessed how frequently samples at five reduced diversity levels would yield survival estimates within the 95% confidence interval (CI) obtained using all 20 clones (Figure 3b). While including 15 clones always produced a survival estimate with the 95% CI assays using only a single clone failed to produce an estimate within the 95% CI over half the time in all three toxicity treatments. Overall, this reinforces the importance of accounting for genetic diversity in toxicity testing, as failing to do so means estimates of reproduction and survival are likely to be highly variable and inaccurate.

**Figure 3.**
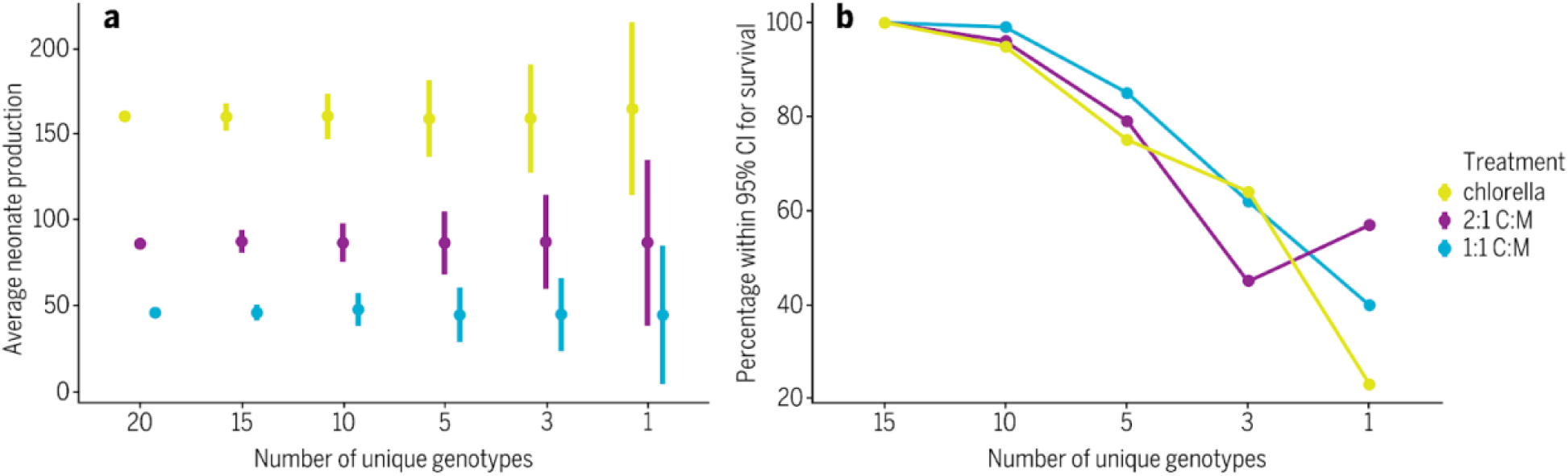
Panel A shows simulations of mean reproductive outputs for experimental designs with varying amounts of genetic diversity of *Daphnia magna* across three treatments of chlorella “C”, 2:1 chlorella:microcystis “2:1 C:M”, and 1:1 chlorella:microcystis “1:1 C:M”. Symbol and error bars represent the mean and standard deviation. A 95% confidence interval was calculated for survival using all 20 clones. Panel B shows simulations of how often estimates of survival fell within this 95% confidence interval when the data were subsetted to include a reduced number of genotypes.

## Discussion

Genetic variation can meaningfully impact population dynamics ^49^, community composition ^50^, ecosystem function ^51^, and underlie rapid adaptation ^10^. Incorporating natural genetic variation in toxicology, which directly informs environmental regulations and policy, is critical to increasing the reliability of benchtop toxicity assays to natural populations. Our data demonstrate that genetic variation, even within a population from a single locality, has a profound impact on toxicological responses in a widely used model system. For example, the moderately toxic diet used in this study could be interpreted as an LC50 (i.e., clone 4) or an LC0 (clone 18 and 19) depending on the clone selected for the assay. When reared on a severely toxic diet, mortality ranged from 0% to 90%. Toxicity studies that do not include intraspecific genetic variation necessarily cannot estimate this variation or provide context on whether the genotype or strain used in the assay is in any way representative of the species or genus. This can lead to considerable uncertainty when extrapolating results from laboratory to field, and have implications for the accuracy and efficacy of risk management strategies ^52^. The ramifications of this oversight are likely to be exacerbated in natural populations where natural selection may strongly shape genetic variation and toxicological responses ^53,54^.

The variation we observed came from a single population of *D. magna* and is doubtless an underestimate of the magnitude of variation that would be found across populations that experience dramatically different exposures to HABs. For example, if the species of focus spans a wide range of ecological conditions, particularly with regards to prior toxicant exposure that may have driven adaptation, there is likely to be substantially more intraspecific variation than we observed within the single population we studied here. Hence, we advocate for the inclusion of multiple genotypes in clonal test organisms or individuals from multiple populations in sexual organisms when conducting toxicity studies. Ideally, studies should survey across a sufficiently wide array of genetic diversity that is similar to populations that could be impacted by the toxicant to allow ‘clone’ or ‘population origin’ to be treated as a random effect in linear mixed models and hence generate reasonable estimates of the variation in key response variables across the populations of interest ^55^.

In practice, we recognize that such robust sampling designs may be challenging and would require shifting effort and focus in many toxicology studies. As a preliminary step, it must be acknowledged by toxicology regulatory agencies that LC50s derived from real, genetically heterogeneous populations will differ at local, regional, or global scale. Each geographic scale presents its own sources of variation and shapes the requirements in terms of the sampling required to achieve the stated objective of a meaningful safety limit recommendation based on an LC50. Therefore, testing multiple genotypes within a single population (local/regional scale) or multiple populations of genotypes (international scale) and reporting a CI of LC50s calculated across each would be a strong step forward. Doing so would provide both a more accurate and robust measure of the LC50, as it would be calculated from multiple genotypes or populations and an indication of the uncertainty. The uncertainty of this estimate would not arise from experimental noise but from real differences in biology. Reporting LC50s would then include both the estimate and the variance around this estimate stemming from genetic variation. This would be useful in discussions about risk and the connection between laboratory assays and toxicological outcomes in natural settings, a key area of growth in translating benchtop experiments to wild populations.

The study here demonstrates that considerable variation is found in toxicological responses within a population. However, it is important to consider the overall scale and scope of variation in toxicological responses. Current methods often examine focal taxa across broad phylogenetic categories (e.g., use of a single surrogate species for a family or order), which is important. But meaningful biological variation can be found across levels of organization. Future work examining variation across biological scales —within populations, across populations, between congeners, and across genera— is critical for parsing the levels of biodiversity most relevant for risk assessment and management action. Identifying the biological scales at which the largest fraction of functional variation is found has been transformative in ecology ^56^ and a similar approach could also lead to a major advance in toxicology.

The level of genomic variation observed across clones in our study demonstrates the large magnitude of genetic variation found within a single population of *Daphnia*. From a toxicological perspective this variation is important both as it provides phenotypic variation, as demonstrated here, and this variation is the raw material on which natural selection can act to drive adaptation. Previous work has suggested that rapid adaptation could be a critical mechanism of suppressing *Microcystis* blooms as field populations of *Daphnia* can rapidly adapt to tolerate *Microcystis* and can suppress the biomass of HABs that contain lethal concentrations of microcystins (3 to 6.5 μg L^−1^) ^43^. These data underscore the need for toxicological theory to integrate genetic diversity, and potentially rapid adaptation in routine testing dose-responses between chronically exposed ^24,42,43,57^ and unexposed ^3,58^ populations might well differ due to genomic variation.

The pace and ecological consequences of resistance evolution has been appreciated for decades ^59^, but a growing body of research is seeking to understand the molecular genetic underpinnings of variation in resistance to toxins ^15,60,61^. We conducted whole genome sequencing of all 20 *D. magna* clones assayed in the study. These data provide a holistic view of the genetic diversity found within a population but do not provide the power required to discover the genetic basis of clonal differences in resistance to *M. aeruginosa*. This is underscored by the lack of associations both between phenotypic variation and genomic differentiation and genomic variation within clones and phenotypic responses. In short, it appears variation in responses to cyanobacteria are not the primary force shaping genetic variation and the raw amount of genetic variation itself is not a key determinant in performance when exposed to toxicants. These results largely fit with the prevailing population genetic theory for genome wide patterns ^62^.

The utility of genomic data in toxicology has been touted as potentially transformative - with a gene-level understanding of resistance presented as a major goal ^63^. It is possible to conduct quantitative trait locus (QTL) mapping or genome wide association studies (GWAS) to find the underlying genomic basis of trait variation. However, generating the resolution needed to identify a causal genomic region, particularly specific genes, would require a herculean effort, a relatively simple genetic basis, and likely some recombinational luck - efforts most likely to yield payoff only in model systems. There is considerable interest in understanding the genetic basis of microcystin resistance in *Daphnia* ^47,48,64^. We investigated the link between variation in genes putatively involved in resistance and a clones’ ability to survive and reproduce when exposed to *Microcystis* and found no significant relationship. Clone 4, which was distinct in both phenotype and genotype for putatively functional loci, does have a private haplotype containing 17 singleton SNPs over a 5 kb window of the tyrosine-protein kinase ABL1. However, this clone had >22,000 unique singleton SNPs making any inference about the function of specific genetic variants exceeding tenuous. Given the complex and environmentally variable genetic architecture of trait variation for many complex traits, we advocate that toxicologists focus genetic studies on assessments of clonal, family, or population level variation. Explicitly including this information on origin in study designs and statistical models will yield powerful insights without requiring major changes in the methods most often employed in the discipline.

## Conclusions

Studies of genetic variation have improved our understanding of ecological and evolutionary processes ^60^. Arguments in favor of incorporating genetic variation, and to an extent evolution, in the environmental risk assessment process are growing stronger as the recognition that meaningful levels of intraspecific genetic variation exist in key toxicological traits ^21,23,52,65–67^. Ignoring genetic variation can lead to inaccurate reporting of LC50 toxicity thresholds ^23^, which may obscure safety measures adhered by companies invested in the commercial production of chemicals and other categories of stakeholders more concerned with toxicants threatening aquatic lifebiodiversity. Here, we show that genotypic and phenotypic differences in *D. magna* tolerance to the aquatic contaminant *M. aeruginosa* were substantial across multiple life-history traits. Significant interactions between growth and reproduction were additionally observed among genotypes in green algae versus cyanobacteria treatments. These results suggest that genetic diversity can strongly influence population dynamics with and without toxicant exposure. Using simulations and bootstrapping approaches we further demonstrate that the common toxicological practice of using a single genotype to calculate toxicity tolerance leads to an estimate outside of the 95% confidence interval >50% of the time. We suggest that integrating intraspecific genetic variation into routine laboratory toxicity studies would improve the reliability and biological realism of LC50 calculations and improve both the environmental risk assessment process and our understanding of the transgenerational impacts of aquatic contaminants on natural populations.

## Methods

### Daphnia magna field collection and culturing

Twenty genotypes of *D. magna* were collected from ‘Langerodevijver’ (LRV; 50° 49’ 42.08’’, 04° 38’ 20.60’’), a small lake (surface area = 140,000 m^2^, max depth = 1 m) within the nature reserve of Doode Bemde, Vlaams-Brabant, Belgium ^68^, in late spring and early summer. LRV has a single basin and has seasonal *Microcystis* HABs typical to many lakes globally and has a large resident population of *D. magna*. Parthenogenetic lines of each genotype were maintained for over five years (approximately 300 generations) in continuous cultures at 20 °C in UV-filtered dechlorinated municipal tap water containing 2 mg C L^−1^ of the green alga *C. vulgaris* (strain CPCC 90; Canadian Phycological Culture Centre, Waterloo, ON, Canada). *C. vulgaris* was grown in COMBO medium ^69^.

To prepare for this study, we isolated one adult female *D. magna* per genotype in separate 50-mL glass tubes inoculated with COMBO medium and *C. vulgaris* at 2 mg C L^−1^, and monitored them daily for reproduction. *D. magna* neonates born within 24 hr were collected from their respective genotypes and individually separated into 50-mL glass tubes as previously described, totalling 10 replicates per genotype and 200 tubes total. These 200 *D. magna* representing 20 genotypes were the founding mothers of this study. All *D. magna* were incubated under constant conditions (temperature of 21 ± 1 °C, cool-white fluorescent light of 600 ± 15 lx, with a photoperiod of 16:8 h light:dark).

### Microcystis aeruginosa culturing

Following our previously described method ^70^, *M. aeruginosa* (strain CPCC 300; Canadian Phycological Culture Centre, Waterloo, ON, Canada) was cultured in BG-11 media and kept in a growth chamber under axenic conditions with a fixed temperature of 21 ± 1 °C, cool-white fluorescent light of 600 ± 15 lx, with a photoperiod of 16:8 h light:dark. The culture was grown for a minimum of one month before preparation for the chronic study. *M. aeruginosa* CPCC 300 produces microcystins-LR (CAS: 101043-37-2, C_49_H_74_N_10_O_12_) and its desmethylated form [D-Asp³]-microcystin-LR (CAS: 120011-66-7, C_48_H_72_N_10_O_12_), which occur widely in freshwater ecosystems ^34,36^ and are toxic to zooplankton.

To prepare *M. aeruginosa* for testing on *D. magna*, an aliquot of the stock was inoculated in 100% COMBO medium for two weeks prior to test initiation and cultured to a cell concentration of 1.25 ± 0.02 x 10^7^ cells mL^−1^. This medium was chosen because it supports the growth of algae and cyanobacteria and is non-toxic to zooplankton ^69^.

### Chronic toxicity study

A 14-d chronic toxicity study was performed in accordance with the test guideline of the Organization for Economic Cooperation and Development to assess the effects of contaminants on the survival, growth, and reproductive output of *D. magna* ^71^. The OECD test guidelines are widely used in toxicity laboratory studies. Three treatments were included in this study: 100% chlorella (non-toxic diet), 2:1 chlorella:microcystis (moderately toxic diet), and 1:1 chlorella:microcystis (severely toxic diet). Following these ratios all *D. magna* were fed 2 mg C L^−1^, corresponding to 3 × 10^6^ cells total, which corroborates with previous literature exposing daphnids to dietary combinations of green algae and cyanobacteria ^41,44,72^.

Following the OECD test guidelines (Test No. 211: Daphnia magna Reproduction Test ^71^), *D. magna* neonates born within 24 hr were collected from their respective genotype, and 10 neonates were individually placed in 50-mL glass tubes containing 50 mL of UV-filtered water and its respective treatment. To ensure there were no systematic differences in ages between *D. magna* neonates that could drive variation in responses within and across clones, a stringent protocol was implemented to remove neonates twice, daily in the morning and at night in preparation for the toxicity test. The UV-filtered water met the validity criteria for the test following an accredited method for *D. magna* culturing and toxicity testing ^73^. A total of 600 animals were used for this study (i.e., 10 replicates per genotype, 200 animals per treatment). Since this was a semi-static test, solutions were renewed 3 × wk on Mondays, Wednesdays, and Fridays by transferring *D. magna* from old to new glass tubes, followed by supplying each *D. magna* with 3 × 10^6^ cells of food (e.g., the 2:1 treatment received 2 × 10^6^ *C. vulgaris* cells and 1 × 10^6^ *M. aeruginosa* cells, corresponding with 2 mg C L^−1^). During this time, survival, growth, reproductive output, and number of broods were recorded to assess for interactions between genotype and treatment effects. Growth (mm) for each replicate was also measured on days 0, 7, and 14. The study was incubated under 400–800 lx cool-white fluorescent light at 20 ± 1 °C with a 16:8 light:dark cycle.

### Genomics sequencing and analysis

Pools of individuals were collected from each clone and DNA was extracted using the QIAGEN MagAttract HMW DNA kit. Extractions were checked for DNA concentration using a Qubit 2.0 fluorometer and libraries for each clone were prepared using the KAPA Hyper Prep kit. The mean DNA concentration and mean total quantity of DNA used for library construction was 6.0 ± 0.4 ng μL^−1^ and 304.3 ± 15.6 ng, respectively. Sequencing of these libraries was generated on a NovaSeq 6000 generating 150bp paired-end reads. Sequences were then trimmed using *trimmomatic/0.39* using a quality threshold of 20 and a min length of 50 and subsequently mapped to the *D. magna* reference genome (GCA_020631705.2) using *bwa/0.7.17* (default parameters). Duplicates were marked using *Picard tools/2.21.2* and then the bam files were input into *freebayes/1.2.0* to generate a VCF of all variant positions among all individuals on the 10 *D. magna* chromosomes (not scaffolds). *Vcftools* was then used to trim to a minimum quality score of 30. This produced a whole genome dataset with an average genome sequencing coverage of 29.5 (min=26.6, max=35.5, SE=0.34) for each of the 20 clones. The total number of variant sites and the number of heterozygous sites for each clone was extracted using *VCFtools.* Genetic distance was calculated using *VCF2dis* (https://github.com/BGI-shenzhen/VCF2Dis). Genotype likelihood (beagle) files were generated using *angsd* (-GL 1 -out D23Genolike -doMaf 2 -SNP_pval 1e-6 -doMajorMinor 1 -doGlf 2), and *pcangsd* was used to generate principal component analysis.

### Water sampling for analysis

Water samples were collected from each treatment at test initiation, solution changes, and termination to quantify the cell concentration, fluorescence, and standard water parameters and were quick-frozen at −80 °C before analysis for microcystins and oligopeptides (Supporting Information, Dataset S1).

### Chemicals and Reagents for water analysis

Water, acetonitrile, methanol, acetic acid, and formic acid were all Optima LC/MS grade solvents and purchased from Fisher Scientific (Tewksbury, MA, USA). Microcystin standards D-Asp^3^-RR, MC-RR, YR, HtyR, LR, D-Asp^3^-LR, HilR, WR, LA, LY, LW, and LF, as well as nodularin and cylindrospermopsin were purchased from Enzo Life Sciences (Farmingdale, NY, USA). Microcystin Dha^7^-LR was purchased from National Research Council Canada (Ottawa, ON, Canada). Microcystin Leu^1^-LR was purchased from GreenWater Laboratories (Palatka, FL, USA). D-Asp^3^-E-Dhb^7^-RR was purchased from Sigma Aldrich. Anabaenopeptin A and B, oscillamide Y, oscillaginin A, microginin 690 methyl ester, aeruginosamide B and C, cyanopeptolin 1040 MB and 1007, and micropeptin 1106 were purchased from MARBIONC (Wilmington, NC, USA). Anabaenopeptin F was synthetically created by the Stockdill laboratory at Wayne State University (Detroit, MI, USA). ^15^N MC-LR was purchased from Cambridge Isotope Laboratories, INC. (Tewksbury, MA, USA).

### Microcystin and cyanopeptide water analysis

Microcystins were measured using a method based on the US EPA method 544 ^74^. Quality assurance and quality control met the criteria set by the US EPA method 544. Dataset S1 depicts the standard curve range, retention time, and linearity, as well as the quantitative and qualitative ions that were used for further identification of the microcystins. Surrogate recovery for the samples were within the US EPA method 544 limit of 60-130%, and the control standards were within the limits of 70-130% set by this same method. All duplicates relative % differences were calculated to be < 30%.

Microcystin and cyanopeptide standards and samples were analyzed using a Thermo Scientific TSQ Altis™ triple quadrupole mass spectrometer (Waltham, MA, USA) with a TriPlus™ RSH EQuan 850 system. Both online concentration analysis and direct inject analysis were completed ^74^. Quantitative results shown use the direct inject analysis method. Detection limits for the online concentration method were between 0.5-5 ng L^−1^, and the standard curve range was between 0.5-500 ng L^−1^.

For direct injection analysis, 10 µL injections of standards and samples were loaded onto a Thermo Hypersil GOLD™ C18 2.1 x 50 mm, 1.9 µm column using a TriPlus RSH autosampler equipped with a 100 µL sampling syringe. The analytical separation gradient consisted of 0.1% formic acid in water (mobile phase A) and 0.1% formic acid in acetonitrile (mobile phase B). At a flow rate of 0.5 mL/min, the gradient started at 24% B and was held for 0.7 min. It was then ramped to 26% B from 0.7–1.7 min, from 26–50% B from 1.7–5.5 min. The column was washed at 98% B from 5.51–6.5 min at a flow rate 1.0 mL/min, then brought back down to original conditions to equilibrate. The total run time was 8.5 min. The column oven temperature was held at 35°C. The standard curve was prepped in LC-MS water and the range measured between 0.05-50 µg L^−1^.

Mass spectrometry analysis was performed using electrospray ionization source (ESI). The MS source settings were as follows: spray voltage 3500 V, ion transfer tube temperature at 325 °C, vaporizer temperature at 350°C, sheath gas 35 (arbitrary units), auxiliary gas 10 (arbitrary units), sweep gas 0 (arbitrary units), collision gas at 1.5 mTorr, and Q1/Q3 resolution set to 0.7 Da.

Control samples and standards were performed every 20 samples during each run. Control standards included negative control (blank) and positive control (10 ppb standard) as well as duplicate and fortified samples. For each standard, a quantitative and qualitative ion transition were chosen for use with selected reaction monitoring (SRM).

### Selective ion mode analysis

For selective ion mode (SIM) analysis, the following masses in Supporting Information, Dataset S1 were investigated because they are known to be produced by *M. aeruginosa* CPCC 300 (Racine et al. 2019). The gradient for the method consisted of 0.1% formic acid in water (A) and 0.1% formic acid in acetonitrile (B). At a flow rate of 0.35 mL min^−1^, the gradient started at 0% B and was held for 1.0 min. It was then ramped to 60% B from 1.0-8.5 min. The column was washed at 98% B from 8.51–10.5 min at a flow rate 0.6 mL min^−1^, then brought back down to original conditions to equilibrate. The total run time was 14 min. The injection volume was 10 µL and the column oven temperature was held at 35 °C. Mass spectrometer source settings were the same as the quantitative method above. The SIM method was accomplished in Q3. The area under the m/z peak was calculated and reported.

### Testing criteria and toxin analysis

Measured microcystin concentrations were 3.3 ± 0.001 μg L^−1^ and 5.1 ± 0.001 μg L^−1^ in the 2:1 and 1:1 toxic treatments, respectively (Dataset S1). Water parameters before and after solution renewals were also within the test criteria (Dataset S1). There were no signs of hypoxia stress on *D. magna* behavior. Following the criteria for survival and reproduction of controls, all test acceptability criteria were met. Mortality and immobilization were first observed within 48 h after microcystin exposure in either treatment, and continued onward throughout the test. This was expected as the microcystin concentrations chosen can be sublethal ^44^ to lethal ^45,46^ in *Daphnia* laboratory assays and are commonly measured in freshwater ecosystems with HABs ^36^.

### Statistical analysis

#### For survival, growth, and neonate production

To test the magnitude of clonal variation, we used a Linear Mixed Effects (LME) model (or a Generalized Linear Model (GLM) for binomial survival data) for each response with clone treated as a random effect and tested for significant differences in fit between models with and without clone included as random effect ^75^. Second, to determine whether the concentration of *Microcystis* influenced the magnitude of clonal variation we conducted Brown-Forsythe tests for homogeneity of variances after generating means for each clone for each response variable. These Brown-Forsythe tests were done sequentially - meaning comparisons were between *Chlorella* and 2:1, then 2:1 and 1:1. Finally, to test for interactions between response variables across *Microcystis* concentrations we used linear models with concentration and clone as fixed effects (*glm* with binomial for survival, *lm* for growth, and *glm* with a gamma distribution for neonate production).

#### Association between clonal genomic diversity, genomic divergence, and phenotypic distance

We tested for an association between the amount of genetic diversity found within clones and clonal survival and reproduction across each of the three diet conditions. To do so we regressed the total number of heterozygous sites and each response variable for each rearing condition. To test for associations between genomic diversity and phenotypic distance, we first generated distance matrices from mean values for all 20 clones. The genomic distance matrix was constructed from all polymorphic sites. The phenotypic distance matrix was constructed using neonate production, time to first brood, total broods, week one growth (day 0-7), week two growth (day 7-14), week one survival, and week two survival for each clone for each diet condition. A two-tailed Mantel test was then conducted to determine whether there was significant correlation between matrices that differed from 0.

We conducted a similar set of analyses exclusively using genes previously identified as involved in microcystin tolerance. These two unique *Daphnia* studies identified potential candidate genes for microcystin tolerance in *Daphnia* through a combination of differential expression analysis (RNAseq) and employing functional annotations ^47,48^. We search the gene names provided by the authors in the NCBI database. If only a single gene region matched the name we included it in our downstream analysis. For gene names that had multiple matches we selected from the most exact. Finally, when authors provided primer regions we used ePCR to identify the gene region that would be amplified in the reference. Using these methods we were able to locate 8 potential candidate loci for genetic variation in microcystin tolerance. We then identified all variant sites across the 20 *Daphnia* clones within these eight candidate gene regions, including 1,000 bp up and downstream of the demarcated gene, for each of 20 clones. We constructed a PCA using this data, conducted a two-tailed Mantel test to determine whether there was significant correlation between differences at these loci and phenotypic variation, and tested for associations between the number of variants in these eight genes and survival and reproduction in high (1:1) *Microcystis* conditions.

#### Examining the effects of assays with reduced diversity

We sought to determine how the choice to conduct lower diversity assays might influence the estimate of the effects of the toxicant on reproduction and survival. For reproductive output we calculated means and standard errors for all 20 clones and for 15, 10, 5, 3, and 1 clonal diversity groups created by subsetting the full dataset 100 times at each diversity level. For survival, we calculated the 95% CI of survival probability using the means of all 20 clones. We then created subsamples containing 15, 10, 5, 3, and 1 clones and determined how frequently they would produce survival estimates within the 95% CI obtained from the full dataset.

All analyses were completed in R version 4.2.2 ^76^.

## Acknowledgements

We thank the Rudman and Fryxell labs for helpful comments and discussions. This work was supported by the Canada First Research Excellence Fund through the Food from Thought Program (JMF), an NSERC Discovery Grant (JMF), the National Institute of General Medicine of the National Institutes of Health (Award #1R35GM147264; SMR), a Banting Postdoctoral Fellowship (SMR), an NSERC Postdoctoral Fellowship (RSS), and a Liber Ero Postdoctoral Fellowship (RSS).

## Supporting Information

Dataset S1: Tables S1-S8, Figure S1

## Author Contributions

R.S.S., S.M.R., L.D.M., and J.M.F. designed the research. R.S.S. and S.M.R. conducted the experiments. J.A.W. and X.W. conducted supporting data collections. S.M.R., R.S.S., and C.I.C. analyzed the data. R.S.S. and S.M.R. wrote the paper with insights from L.D.M., C.I.C., and J.M.F. All authors read, amended, and approved the final manuscript.

## Competing Interest Statement

The authors declare no competing interests.

## Data Availability Statement

The primary dataset that support the findings of this study are included in the Supporting Information.

## Notes

### Competing Interest Statement

The authors have declared no competing interest.

### Summary of Updates

The Main Text has been revised; new panel created in Figure 1; author affiliations updated; Supplemental files updated.

